# Molecular Modeling Suggests Homologous 2-APB Binding Sites in Connexins 26 and 32

**DOI:** 10.1101/287326

**Authors:** Thomas W. Comollo

## Abstract

Connexins are the transmembrane pore forming proteins that participate in gap junctions; connections between cells that certain nutrients and other molecules can pass through. There are several kinds of connexins (Cx) named based on their weight in kilodaltons. 2-aminoethoxydiphenyl borate (2-APB) is a small molecule inhibitor (SMI) of Cx26 and Cx32. Knock out of Cx32 and also blockage of Cx32 by 2-aminoethoxydiphenyl (2-APB) has been suspected to be beneficial in not only, drug induced liver toxicity, but also in blocking the propagation of an inflammatory signal. If the binding site of 2-APB can be determined, virtual screening for additional, perhaps more specific Cx26 and Cx32 blocking SMI can be carried out. Our modeling suggests that 2-APB binds to similar sites inside the pores of Cx26 and Cx32. Here 2-APB interacts with the conserved ILE82 and THR86. These residues hold the same numbering in both CX26 and CX32. This suggests these residues have a high level of conservation and importance Further virtual screening results imply molecules with similar activity on Cx26 and Cx32 as 2-APB can be found.

**Background:** 2-APB has been shown to block Cx26 and Cx32.

**Results:** Docking of 2-APB to Cx26 and CX32 finds conserved binding site in pore. Suggestive that 2-APB has conserved homologous binding site on Cx26 and Cx32.

**Conclusion:** Mutational studies of 2APB and ILE82 and THR86 in Cx26 and Cx32 appear warranted. Additional virtual screening could yield 2-APB analogs that act on Cx26 and Cx32.

**Significance:** Potential for developing gap junction blocking compounds.

## INTRODUCTION

Connexins (Cx) are a family of vertebrate gap junction forming proteins named based on their weight in kDa. They exist as homo or hetero hexamers that can exist alone as a hemi channel or two bridging two cells together as a gap junction [1]. 2-aminoethoxydiphenyl borate (2-APB) is a compound initially shown to trigger IP_3_ Ca^2+^ release in cerebellar microsomes [2]. 2-APB and analogs have been demonstrated as small molecule inhibitors (SMI) to block Cx hemichannels [3, 4]. Small molecule inhibitors (SMI) that block gap junctions in liver, namely 2-APB, were shown to block communication at the hepatic gap junction and deter drug induced liver toxicity.

Our modeling suggests conserved residues ILE82 and THR86 are part of a 2-APB interaction site inside the pore of CX26 and CX32. Our results also suggest that it may be possible to find other Cx SMI through virtual screening.

## EXPERIMENTAL PROCEDURES

### Molecular Modeling

2-APB’s docking to human Cx26 and Cx32 using AutoDock Vina [5] was made possible by using a dummy carbon atom in the place of the molecule’s boron. The 2apb model was sketched, hydrogen added, and was then minimized in SYBYL-X [6]. Gasteiger-Huckel charges were then calculated. A note was made of these charges. A. pdbqt was then created of the model in AutoDock Tools [7], making sure not to change the hydrogen layout. This. pdbqt was then hand edited to add the previously SYBYL-X calculated charges to the appropriate atom lines in the 2apb. pdbqt.

Cx26’s structure (pdb 2zw3) [8] was converted to a. pdbqt using AutoDock Tools. The Cx32 model was created from the provided alignment (see supplementary material) of the sequence of pdb 2zw3 to that of Cx32 as obtained from the Uniprot data base [9]. This alignment was performed in Clustal-X, as distributed with VegaZZ and used defaults, save for a gap open penalty changed to a value of 20. The homology model of a single Cx32 subunit was then created using the Swiss Model server in alignment mode [10]. The multimer of the Cx32 hemi channel was then assembled using locally run M-ZDock [11]

The The 2-APB model was docked to the Cx26 and the Cx32 models, in search areas encompassing the complex’s pore. The virtual screening was run using the same search areas. Ligand set up and AutoDock Vina were run in “for each” loops (procedure at http://home.fatsilicodatapharm.com/home/virtual-screening-methodology). The best scoring dockings were inspected for conformations in the same ILE82, THR86 docking pocket as the 2-APB model.

## RESULTS

### 2-APB docked to a homologous, conserved pocket in the pore of Cx26 and Cx32

Docking a model of the SMI, 2-APB (Figure 1), to the entire pore region of a Cx26 crystal structure and a Cx32 homology model found the best scoring poses for each docking were similar, in a homologous, conserved pocket in the pore of both macromolecules (see Figure 2). The best score for 2-APB docked to Cx-26 was −7.6kcal/mol and for Cx32 it was −8.2 kcal/mol. In the case of both Cx26 and Cx32, the 2-APB docked model was wedged between the first alpha helix and those that form the walls of the pore. The 2-APB docked model closely contacted ILE82 and THR86. The proposed binding site is primarily contained within a single one of each of the six identical subunits that form the pore. This is our pocket of interest (POI).

**Figure 1.**
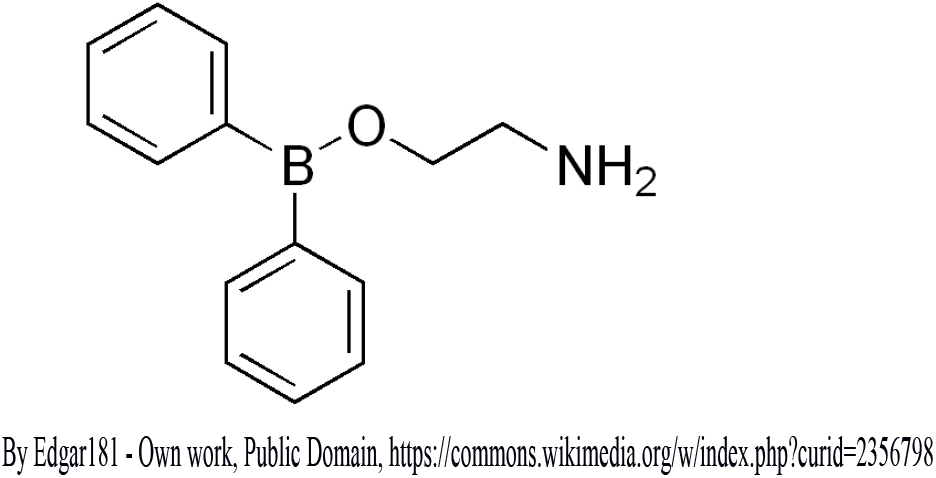
Structure of *2-aminoethoxydiphenyl borate (2-APB)*. 2-APB has been shown to block CX26 and CX-32 gap junctions.

**Figure 2.**
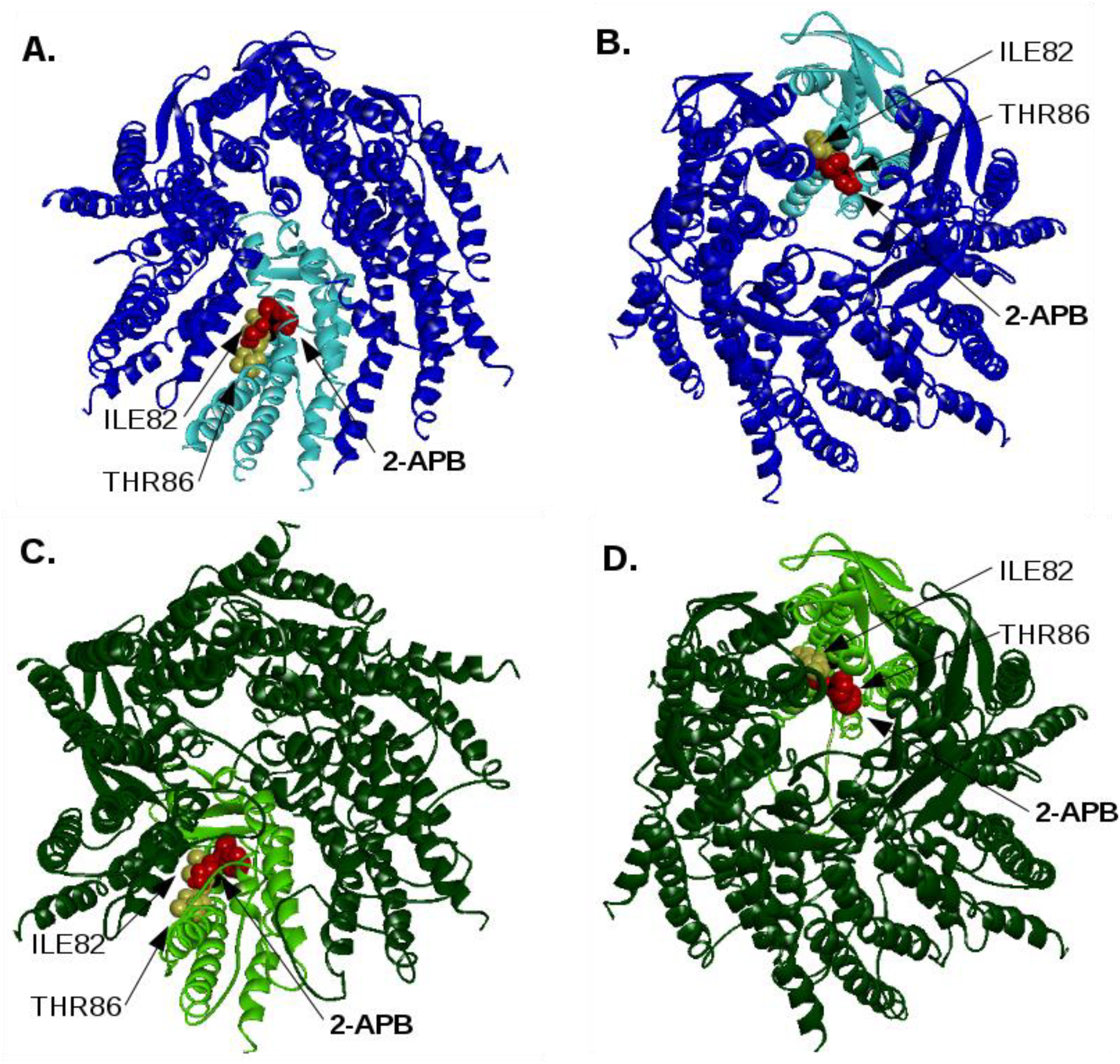
Docking of 2-APB model to pore of CX-26 crystal structure and CX-32 homology model show homologous potential binding sites. 2-APB contacts ILE82CX26 and THR86CX26 in CX-26 docking and the homologous ILE82CX32 and THR86CX32 in the CX-32 homology model docking model. A) Intracellualar view of 2apb docked to CX-26 hemichannel (PDB entry 2zw3). B) View from extracellular side of of membrane looking at docking of 2-APB to CX-26 hemichannel. View from intracellular of 2-APB docked to CX-32 homology model built from sequence alignment to PDB 2zw3. Extracellular view of same docking model.

**Figure 3.**
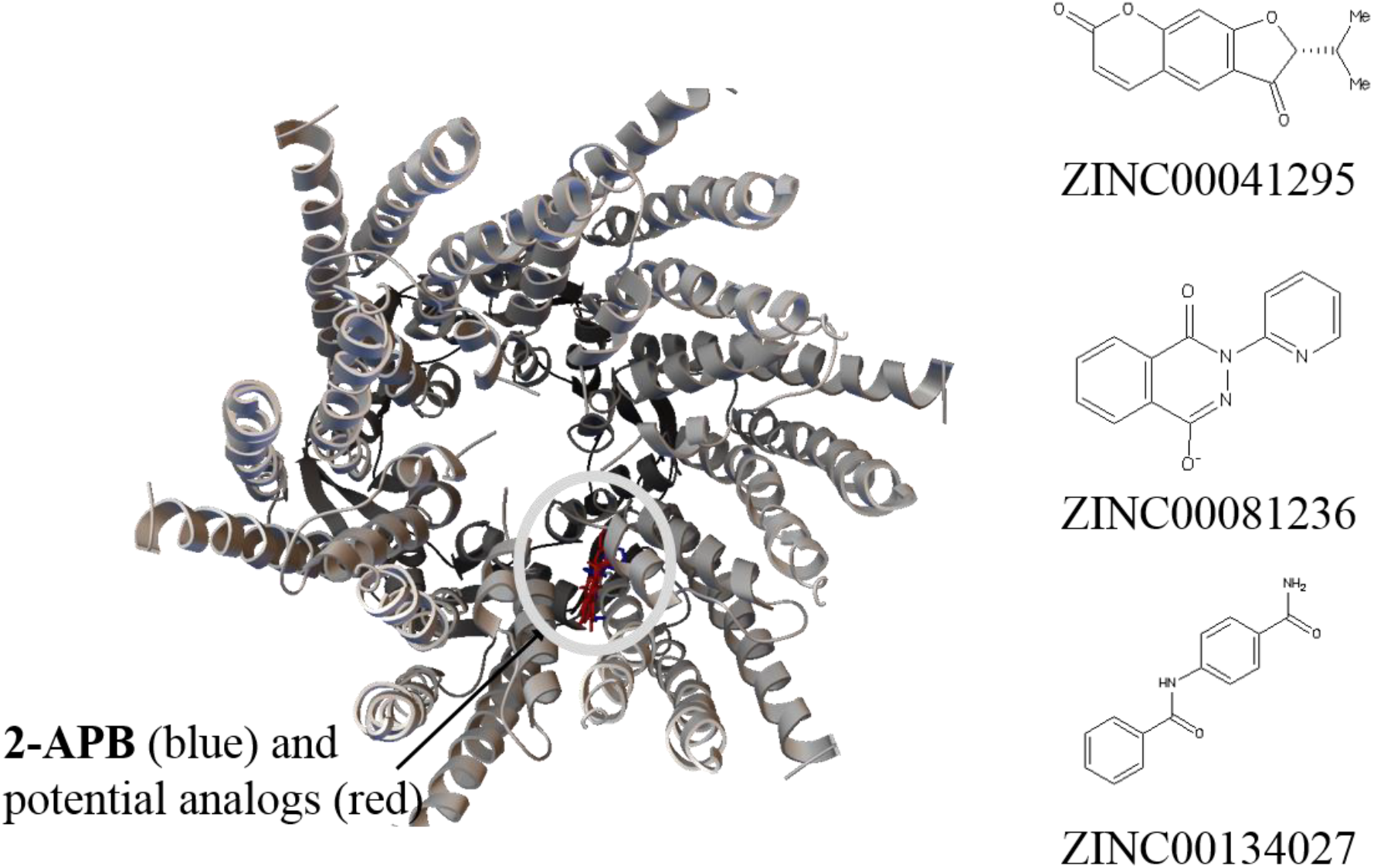
Virtual screening to POI yields theoretical “hits”. A short virtual of ~3,000 fragment size compounds to pore of CX-26 crystal structure yields multiple compounds that similarly fit the possible 2-APB site with better docking scores than our 2-APB model. Some of these are shown docked above in red stick and 2-APB in blue stick. Their structure and ZINC (zinc.docking.org) numbers are at right.

### Virtual screening predicts other compounds will bind Cx26 2-APB binding site

Virtual screening of ~3,000 compounds from the zinc.docking.org data bank found several that were predicted to bind the Cx26 POI with scores better than 2-APB’s −7.6 kcal/mol. ZINC 00041295 ((2R)-2-isopropylfuro[3,2-g]chromene-3,7-quinone-8.3) docked with a best scoring conformation of −8.6 kcal/mol. ZINC00081236 (LS-109167) and ZINC00134027 docked in the POI with best conformation scores of −8.4 kcal/mol and −8.0 kcal/mol respectively.

## DISCUSSION

The Cx26 and Cx32 pore binding site for 2-APB appears to be conserved. They are highly homologous proteins that are both blocked by the SMI, 2-APB. To our knowledge, the binding site for 2-APB on Cx26 and Cx32 has not been published. ILE82 and THR86 hold such a high level of conservation that they have the same numbering in both Cx26 and Cx32. This lends value to this prediction. It seems likely that these residues that hold so much in common between Cx26 and Cx32 would have a similar and important function. Homologous proteins, both inhibited by the same SMI would likely have similar sites of interaction.

If this POI is accurate then further virtual screenings can be carried out for molecules with 2-APB like blocking effects on connexins, but perhaps with more specificity. The two interacting residues were the same between the two, but a fully drug size molecule may be able to exploit differences in the macromolecules around ILE82 and THR86. Further, there may be as many as six of these sites on each Cx26 and Cx32 hemichannel. It may be useful to determine the exact stoicheiometry of 2-APB to Cx subunit for effective block. Is one 2-APB molecule sufficient to block a hemichannel or are more, maybe six required?

The accuracy of these predictions are not verified in the experimental laboratory. This is required if these predictions are to be held true. Molecular modeling can be a powerful tool for hypothesis generation and explanation, but is highly subject to delivering inaccuracies, even when performed correctly. If these predictions are true, however, other 2-APB like Cx26 and/or Cx32 SMI, with possible uses such as treating drug induced liver toxicity [12], may only be a few screenings away.

*Thank you to Michael Salling, Ph.D. for helpful comments*.

