## Supplementary Materials for "Molecular Modeling Suggests Homologous 2-APB Binding Sites in Connexins 26 and 32"

### CLUSTAL X (1.83) MULTIPLE SEQUENCE ALIGNMENT

#### File: C:Documents and SettingsThomas W. ComolloDesktop -23-10connexin-32_2.ps  Date: Sat Jul 24 00:55:09 2010
#### Page 1 of 1

```
                           *:*  * *:*.***:***:**::**:*:******:*****:.*****::.*:********:.****::*****:***:****:*************:::* :* *::: * :.:   .:**:* :**:*.*:*****. *:.**::***.*
W3_A|PDBID|CHAIN|SEQUENCE MDWGTLQTILGGVNKHSTSIGKIWLTVLFIFRIMILVVAAKEVWGDEQADFVCNTLQPGCKNVCYDHYFPISHIRLWALQLIFVSTPALLVAMHVAYRRHEKKRKFIKGEIKSEFKDIEEIKTQKVRIEGSLWWTYTSSIFFRVIFEAAF   150
    sp|P08034|CXB1_HUMAN MNWTGLYTLLSGVNRHSTAIGRVWLSVIFIFRIMVLVVAAESVWGDEKSSFICNTLQPGCNSVCYDQFFPISHVRLWSLQLILVSTPALLVAMHVAHQQHIEK-KMLRLEGHGDPLHLEEVKRHKVHISGTLWWTYVISVVFRLLFEAVF   149
                    ruler 1.......10........20........30........40........50........60........70........80........90........100.......110.......120.......130.......140.......150
```

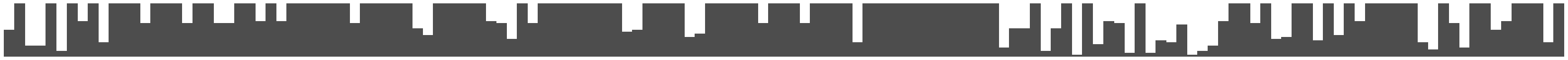

```
                           *****::* *::* *****:.:***********************:*.*****:***:*: **:** *: ::::
W3_A|PDBID|CHAIN|SEQUENCE MYVFYVMYDGFSMQRLVKCNAWPCPNTVDCFVSRPTEKTVFTVFMIAVSGICILLNVTELCYLLIRYCSGKSKKPV----------------------------------------------------------   226
    sp|P08034|CXB1_HUMAN MYVFYLLYPGYAMVRLVKCDVYPCPNTVDCFVSRPTEKTVFTVFMLAASGICIILNVAEVVYLIIRACARRAQRRSNPPSRKGSGFGHRLSPEYKQNEINKLLSEQDGSLKDILRRSPGTGAGLAEKSDRCSAC   283
                    ruler .......160.......170.......180.......190.......200.......210.......220.......230.......240.......250.......260.......270.......280....
```

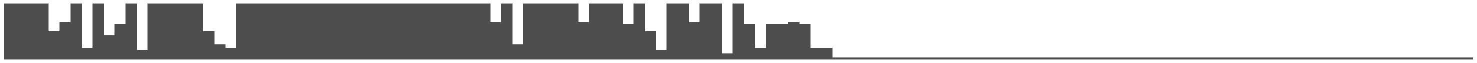
